# No decoy effect in bees: rewardless flowers do not increase bumblebees’ preference for neighbouring flowers

**DOI:** 10.1101/2025.03.24.644861

**Authors:** Mélissa Armand, Leonhard Herrnberger, Clara Jung, Tomer J. Czaczkes

## Abstract

Nectarless flowers are common among flowering plants, which often retain colour-changed, rewardless flowers instead of shedding them. Yet, how these flowers influence pollinators’ foraging choices within an inflorescence remains unclear. We hypothesised that rewardless flowers in an inflorescence may act as “decoys”, causing the rewarding flowers in the inflorescence to be perceived as more valuable by contrast. Using artificial inflorescences, we presented individual bumblebees (*Bombus terrestris*) with a binary choice between two equally rewarding inflorescences, one of which included additional unrewarded, differently coloured flowers. We found that the presence of rewardless flowers did not increase bees’ preference for neighbouring flowers, nor did it affect their overall choice between inflorescences. However, bees quickly learned to avoid the unrewarded flowers, drastically reducing visits and probing within a few foraging bouts. We review research on decoy effects in bees, and find very little support for their presence. Our findings contribute to the growing body of evidence that rewardless flowers do not induce decoy effects in bees, and highlight the need for further research into the ecological role of nectarless flowers within floral patches. It may be time to abandon the search for classical decoy effects in pollinators.

## INTRODUCTION

Decision-making in animals is context-dependent: preferences arise from comparisons between available options, rather than from their only intrinsic value (Tversky, 1969; Tversky and Simonson, 1993; Rosati and Stevens; 2009; Owen et al., 2017). For example, a food reward may seem more appealing next to a poorer option, or a potentially acceptable mate may become irrelevant if a better one is available (Bateson et al., 2003; Wendt et al., 2019; Poissonnier et al., 2024). Such comparative evaluation can sometimes lead to seemingly “irrational” choices, where the chosen option is not objectively superior but appears favourable in context (Ariely, 2009).

One prominent example of this is the decoy effect, a cognitive bias where adding a clearly irrelevant, inferior option to a choice set shifts preferences among the remaining options (Huber et al., 1982). This phenomenon has been widely observed in both real-world contexts and experimental settings, affecting humans (Simonson and Tversky, 1992; Ariely, 2009; Marini et al., 2020) as well as non-human animals across diverse taxa, including mammals (Scarpi, 2011; Parrish et al., 2015; Jackson and Roberts, 2021), invertebrates (Shafir et al., 2002; Sasaki and Pratt, 2011; Hemingway et al., 2024) and even an unicellular organism (Latty and Beekman; 2011).

Flower-visiting insects navigate a multitude of floral options which differ in various traits that can be compared — a context particularly relevant to decoy effects, as reviewed by Latty and Trueblood (2020). Notably, many flowers within floral patches are naturally nectarless (Thakar et al., 2003). The persistence of such unrewarded flowers in plant populations remains unclear, as well as their potential impact on pollinator behaviour. Could these rewardless, irrelevant flowers act as decoys and influence the foraging decisions of pollinators?

Most decoy effects create an attraction effect, where the presence of an irrelevant decoy makes the option that most resembles it more appealing, shifting attention away from other options (Huber et al., 1982). The best studied example of this is the asymmetrically dominated (AD) decoy: clearly worse than a target option in one attribute but comparable to competitor options, making the target stand out as the best choice (Huber et al., 1982; Huber and Puto, 1983; Wedell, 1991). AD decoys have been shown in the context of foraging choices in pollinators: in honeybees, a decoy flower shifted preference towards flowers with deeper corollas over more concentrated nectar (Shafir et al., 2002) or towards flowers with a higher sucrose concentration over flowers offering warmer nectar (Tan et al., 2014b). In bumblebees, decoy effects were tested along two attributes, nectar concentration and reward rate, but an effect was only observed for reward rate (Hemingway et al., 2024).

Another class of decoys are phantom decoys: they create an attraction effect by introducing an option that appears desirable but is unavailable at the time of choice (Pratkanis and Farquhar, 1992; Trueblood and Pettibone, 2017). This unavailability leads decision-makers to favour the most similar alternative to their preferred, “sold out” option. Nectarless flowers in nature, for instance, may act as phantom decoys (Latty and Trueblood, 2020). However, few studies have examined the effect of empty flowers on pollinators in the context of a decoy effect, and findings are inconsistent. In honeybees, Tan et al. (2014a) found that an attractive but unavailable feeder increased preference for the most similar available option, whereas an unattractive phantom decoy had no effect. Conversely, Forster et al. (2023a) found that introducing an empty, previously rewarding flower did not shift bees’ preferences between the available flowers, and a similar lack of effect was observed in bumblebees (Forster et al., 2025).

In nature, pollinators tend to avoid unrewarding flowers (Cresswell, 1999; Smithson and Gigord, 2003) and spend less time on inflorescences containing empty flowers (Biernaskie et al., 2002; Hirabayashi et al., 2006). Rewardless flowers also increase bee movement between flowers and often lead to higher patch abandonment (Cresswell, 1990; Ishii et al., 2008; Nakamura and Kudo, 2016; Forster et al., 2023a). Yet, their influence on pollinator behaviour is ambiguous. Bees may prefer revisiting a known unrewarding flower over exploring an unfamiliar flower colour (Dyer & Murphy, 2009) and may be drawn to flowers with similar colours or attributes if they were previously rewarding (Internicola et al., 2007; Tan et al., 2015). Conversely, encountering empty flowers might instead push bees to seek out flowers with distinctly different colours (Smithson and Macnair, 1997; Smithson and Gigord, 2003).

Rewardless flowers are widespread among flowering plants (Thakar et al., 2003; Smithson and Gigord, 2003). Beyond the energetic cost of nectar production (Bell, 1986; Pyke, 1991), maintaining empty flowers may help reduce visitation of foragers to inflorescences, thereby limiting self-pollination between flowers (Thomson and Plowright, 1980; de Jong et al., 1993; Biernaskie et al., 2002). Among rewardless flowers, some never produce nectar (Gilbert et al., 1991; Gaskett, 2011), while others gradually decrease nectar production as they age (Weiss, 1991; Gilbert et al., 1991). Aging or pollinated flowers often change colour (Thakar et al., 2003; Brito et al., 2015) even within an inflorescence (see **Fig. 1A**), potentially signalling pollinators which flowers are unproductive (Weiss, 1991). Here, we propose an alternative hypothesis: that differentially coloured, unrewarding flowers may act as decoys, which by contrast causes the nearby productive flowers to be perceived as more valuable when compared to those in competing inflorescences. Successive contrast effects should also contribute to an increased preference for a rewarded flower experienced directly after experiencing an unrewarded flower (Bitterman, 1976; Wendt et al., 2019).

**Figure 1:**
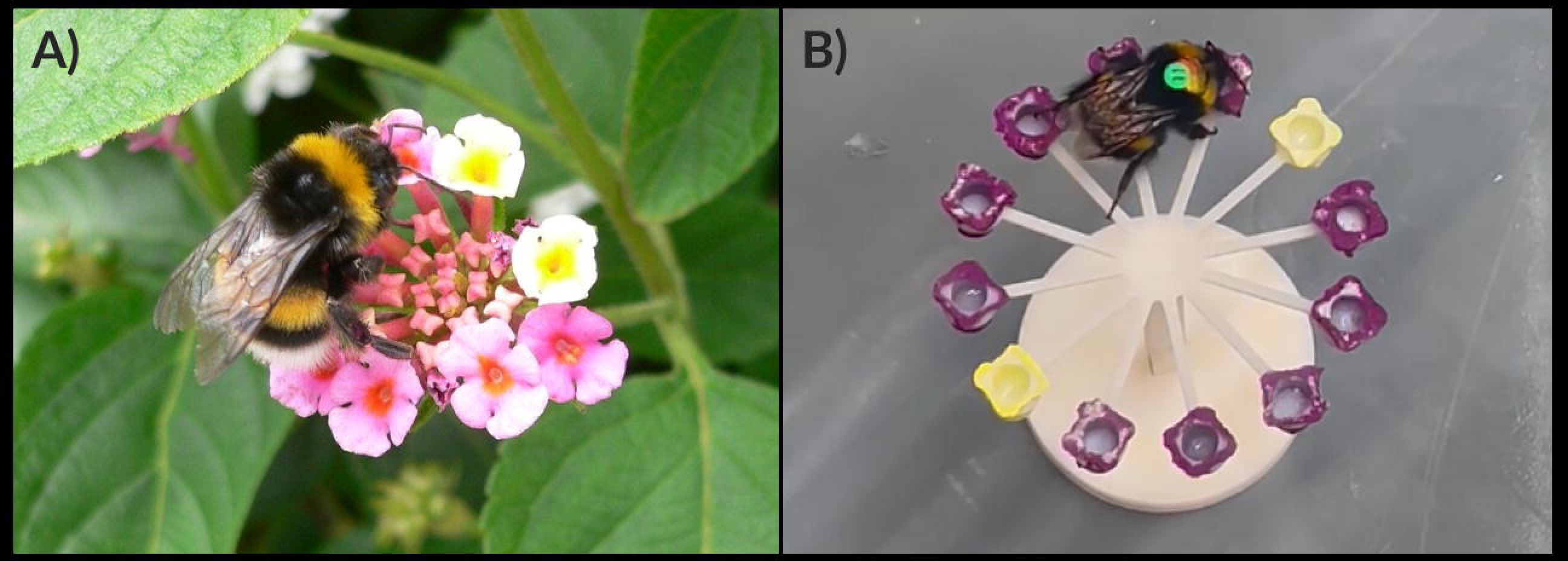
Bumblebee foraging on an inflorescence with differentially coloured flowers. **A)** Inflorescence of *Lantana camara*: newly opened, fertile flowers are yellow, while older, previously visited flowers turn orange to magenta due to anthocyanin accumulation triggered by pollination (Ram & Mathur, 1984). Photo ©Luigi Strano. **B)** Artificial inflorescence used in experiment: purple flowers provided high-quality sucrose solution (50% w/w), while yellow flowers were rewardless and filled with plain water.

In this study, we investigated whether the presence of unrewarded flowers influenced the preference of bumblebees (*B. terrestris*) between two inflorescence options. To test this, we compared bees’ overall preference between two equally rewarding inflorescences, one blue and one purple. In the control group, both inflorescences contained only rewarding flowers, whereas in the decoy groups, one inflorescence included two additional yellow, unrewarded flowers. We hypothesized that these differently coloured, rewardless flowers would act as decoys, enhancing the perceived value of neighbouring rewarding flowers by contrast. Therefore, we predicted that: (1) within groups, bees would prefer rewarding flowers in the inflorescence with decoy flowers, and (2) across groups, bees would have an increased preference for a given inflorescence (blue or purple) when it contained decoys, compared to when it did not.

## MATERIAL AND METHODS

### Colony setup

Commercial *Bombus terrestris* colonies were purchased from Koppert (The Netherlands) and kept under controlled laboratory conditions at 22–24°C with a 14:10 light:dark cycle. Colonies were housed in plastic nestboxes (23 × 21 × 12 cm), modified from their original transport containers with a removable Plexiglas lid for easy handling of the bees. Each nestbox was connected to its respective flight arena (60 × 35 × 25 cm) via a transparent tube leading to a small chamber (6 × 5 × 3 cm). The chamber featured a second tube that provided direct access to the arena and was fitted with transparent, removable shutters to regulate bee movement between the nest and arena (see apparatus **Fig. 2**).

**Figure 2:**
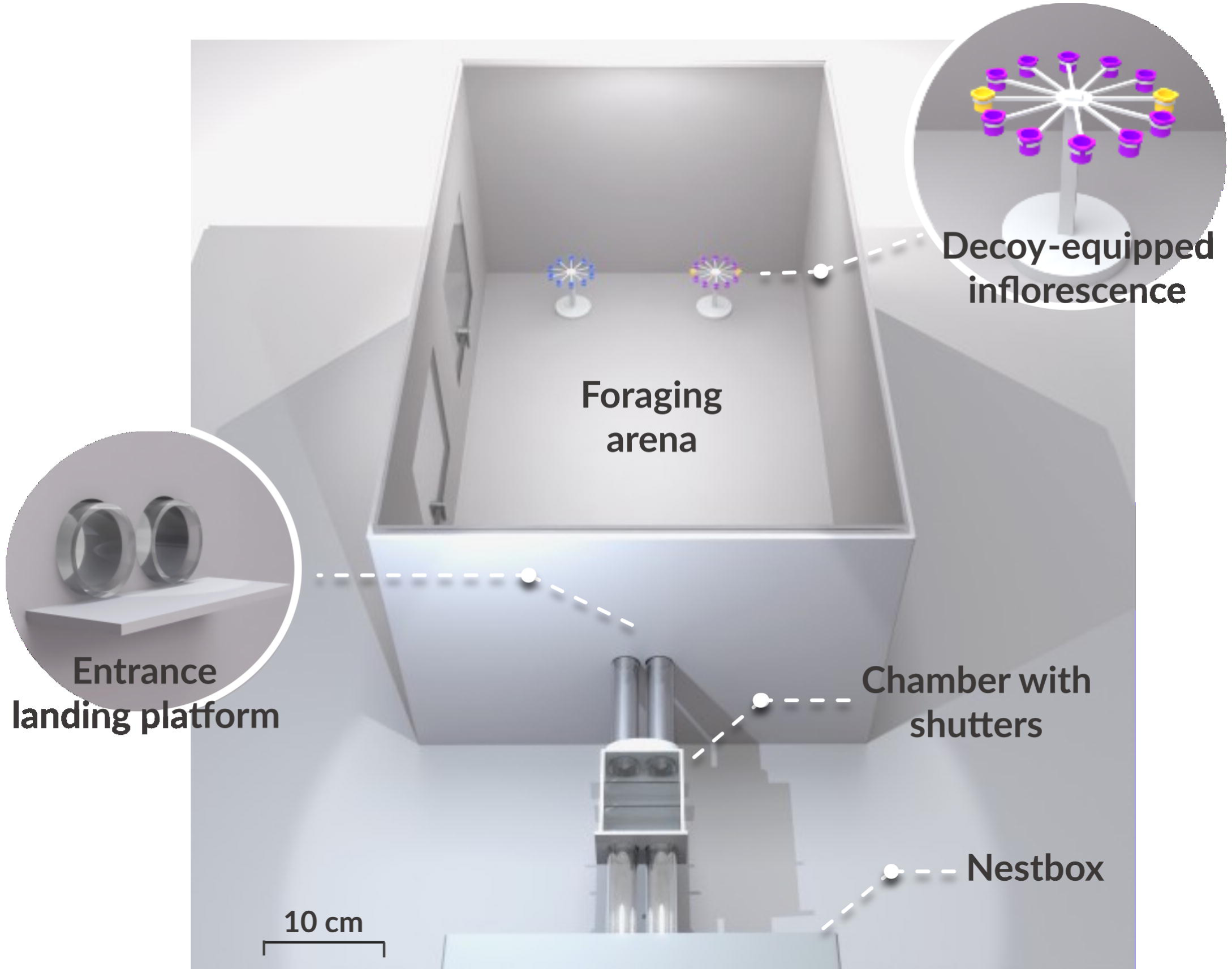
Top view of the experimental setup, to scale. The 3D model, created in Blender, illustrates a training bout featuring a purple, decoy-equipped inflorescence positioned to the right and a blue inflorescence to the left, as oriented from the bee’s perspective upon entering the arena.

Bees were provided daily with pollen balls made from a mixture of organic flower pollen pellets and 35% (w/w) sucrose solution, placed directly in the nestboxes. During the day, workers foraged freely on artificial inflorescences in the flight arena, which offered 35% (w/w) sucrose solution and were regularly refilled. A total of 92 bees participated in the experiment: 10 bees from three colonies in February 2023 and 82 bees from five colonies between October and November 2023 (see colony details in Supplement **S1**).

### Artificial inflorescences

Bees foraged on artificial inflorescences mimicking an umbel, composed of small flowers symmetrically arranged in a circular pattern radiating from a central point (see **Fig. 1B**). Each flower consisted of an opaque cup (0.4 cm diameter, 0.6 cm height) filled with 35% (w/w) sucrose solution. The inflorescences were 3D-designed and printed using white resin and came in two sizes: 10-flower inflorescences (3.4 cm diameter, 5 cm height) and 12-flower inflorescences (4 cm diameter, 5 cm height). In the experiment, the 12-flower inflorescences included two additional unrewarded “decoy” flowers filled with plain water.

Flowers on all inflorescences were spaced 1 cm apart. To collect sucrose, the bees had to fly to an inflorescence, land, and extract the solution from the cups. Flowers were painted blue (peak reflectance ∼470 nm), purple (peaks ∼420 and ∼670 nm), or yellow (weak peak from ∼530–700 nm, see Supplement **S2** for reflectance curves). In the experiment, inflorescences were either all blue or all purple, with the decoy-equipped inflorescence containing two additional yellow decoy flowers. Blue and purple were used as colour cues for rewarding flowers because naïve bees naturally prefer them (Lunau, 1990; Gumbert, 2000), while yellow was chosen as a contrasting colour to clearly differentiate the decoy flowers from the neighbouring blue or purple flowers in the decoy inflorescences. Between experimental sessions, inflorescences were left unpainted in neutral white to prevent bees from associating experimental colours with rewards (see below).

### Pre-training

Between experimental sessions, conducted in the afternoons from 1 to 4 pm, bees were allowed free access to the flight arena to familiarise themselves with the artificial inflorescences. The arena contained four all white inflorescences — two 10-flower and two 12-flower inflorescences — randomly positioned, to ensure bees were equally exposed to both sizes. Individual flowers were regularly replenished by an experimenter using a pipette through the arena’s side doors. Bees observed foraging regularly on the inflorescences were captured and marked on the thorax with uniquely numbered, coloured tags, and considered for selection in the experiment on the same day.

### Training

Bees remaining in the arena from pre-training were returned to the nestbox using forceps, the arena was wiped cleaned with 70% ethanol, and access was closed. When a tagged bee entered the chamber leading to the arena recording began, and training started. Bees were assigned to one of two treatments:

1. **Control treatment** (n = 31 bees): Bees were presented with two 10-flower inflorescences, one with all purple flowers and the other with all blue flowers. Each flower offered 4 μL of high-quality sucrose solution (50% w/w). This treatment tested the bees’ relative preference between the purple and blue inflorescences, both of which offered the same total amount of sucrose.
2. **Decoy treatments** (n = 61 bees): Bees were presented with a 10-flower inflorescence (all purple or all blue) and a 12-flower inflorescence with 10 flowers of the other colour (purple or blue) plus two additional yellow, decoy flowers. Each flower contained 4 μL of high-quality sucrose solution (50% w/w), except the decoy flowers which were filled with 4 μL of plain water, effectively making both inflorescences equally rewarding. For 32 bees, the decoy-equipped inflorescence was blue (“Blue decoy” treatment) and for 29 bees, the reverse (“Purple decoy” treatment). These treatments tested whether the presence of decoy flowers influenced bees’ relative preference between inflorescences compared to the control treatment.

Preferences were assessed at the population level by comparing treatment groups (control vs. decoys) rather than testing individual bees before and after introducing decoys. This approach avoided repeated trials with the same bees, eliminating the potential influence of prior exposure and learning effects.

The inflorescences were positioned at the centre of the arena, one on the left and one on the right, spaced 20 cm apart and 60 cm from the entrance. In the decoy treatments, decoy flowers were consistently placed on opposite sides within each inflorescence (left and right) to ensure their colour was clearly visible to bees entering the arena (see **Fig. 2**). Each bee was randomly assigned a colour and side for each inflorescence type, and these assignments were kept consistent across all bouts to help bees associate each inflorescence with visual and location cues.

In each foraging bout, both inflorescences provided a total of 80 μL of sucrose solution, which is below the average crop capacity of bumblebees (120–180 μL; Lihoreau et al., 2010), ensuring that all rewards could be collected. Inflorescences were replaced with clean ones each new bout to prevent scent marks from influencing subsequent foraging (Goulson et al., 2000; Saleh et al., 2007). Bees were required to probe at least one decoy flower (i.e., extend their proboscis into the cup) during the first bout to continue in the experiment.

Video analysis revealed that bees systematically walked on the surface of the inflorescence between neighbouring flowers, collecting them sequentially before switching to the second inflorescence, after fully depleting the first (see video example in Supplement **S3**). To ensure bees associated yellow flowers with the absence of reward, the decoys were filled with plain water rather than left empty. Pilot trials showed that empty decoys were treated as already-depleted flowers, with bees walking over them without probing, which could weaken the decoys’ intended effect.

### Binary choice test

After completing all six foraging bouts, each bee performed a final test bout with two inflorescences (purple vs. blue), both unrewarded (plain water) and without decoys. The positions of the inflorescences were kept consistent with earlier bouts to help the bees anticipate each inflorescence location and minimise rushed or random choices. We recorded the first inflorescence choice as an indicator of its preference.

### Data analysis

Each foraging bout was video-recorded by a camera (Sony HDR-CX220) positioned above the flight arena. Bee behaviour was analysed using the event-logging software BORIS (v8.6). For each bee, we recorded in every foraging bout: (1) the first inflorescence visited (blue or purple; decoy-equipped or not), (2) the first flower landed on within the inflorescence (blue, purple, or yellow), and (3) whether a decoy flower was probed (i.e., bee’s proboscis extended into the cup).

We tested the hypothesis that the presence of decoy flowers would change bees’ relative preference between the blue and purple flowers, by comparing (1) the overall preference for each inflorescence in the control treatment to those in the decoy treatments, and (2) the preference for each inflorescence within treatments. We built a generalized linear mixed model (GLMM) with the first inflorescence choice (blue or purple) in the binary test as the response variable, and treatment (“control”, “blue decoy” or “purple decoy”) and inflorescence side (“left” or “right”) as predictors. Random effects included bee colony and experimental session, and the model was fitted using a binomial distribution. We examined whether the estimated proportions of first inflorescence choice differed between treatments using post hoc pairwise comparisons with a Tukey correction. Additionally, we tested whether the estimated proportions within each treatment differed from a 0.5 probability using a post hoc Wald *z*–test.

We then tested whether probing a decoy flower increased the likelihood of bees first visiting the decoy-equipped inflorescence in the next bout, favouring nearby rewarded flowers over those of the other inflorescence. We used a GLMM with a binomial distribution, with inflorescence choice (1 for the decoy-equipped inflorescence, 0 otherwise) as the response variable. The fixed effects included whether a decoy was probed in the previous bout (1/0), the foraging bout, and their interaction. Random effects included individual bees nested within their colony and experimental session. The binary test bout and all training bouts were included in the analysis except the initial bout, where decoys could not have been previously probed. Post hoc pairwise comparisons with a Tukey correction tested differences between conditions (decoy probed or not in the previous bout).

Lastly, we examined whether the frequency of decoy flowers probed varied across bouts during training. We built a GLMM with a binomial distribution, with decoy probings as the binary response variable (1 for probing, 0 otherwise), foraging bout as a fixed effect, and individual bees nested within their colony and experimental session as random effects. All training bouts were included in analysis except the first, as bees retained in the experiment were required to probe a decoy flower during the first bout. We performed Post hoc pairwise comparisons with a Tukey correction to test if the estimated proportions differed between bouts.

Data processing was performed with Python (v3.11, Python Software Foundation, 2023) using the libraries *pandas* (McKinney, 2010) for data structuring, *seaborn* (Waskom, 2021) and *Matplotlib* (Hunter, 2007) for data visualisation. Statistical analyses were conducted in R (v4.1, R Core Team 2022) using the *glmmTMB* package (Brooks et al., 2017) for GLMMs, and *emmeans* (Lenth, 2020) for post hoc tests. Model residuals were evaluated with the *DHARMa* package (Hartig, 2020). Complete statistical analyses and datasets are available on Zenodo (https://doi.org/10.5281/zenodo.14995342).

## RESULTS

Overall, 92 bumblebees were tested across all treatments, with 61 bees in the decoy treatments. We analysed 363 first visits from training bouts 2–6 and the binary test bout.

### Effect of decoy flowers on floral preferences

The presence of unrewarded flowers had no impact on bees’ preference between the blue and purple inflorescences (GLMM, binomial family; treatment: X² = 1.23, df = 2, *p* = 0.54), and no effect of inflorescence side was observed (side: X² = 0.0006, df = 1, *p* = 0.98). The likelihood of first visiting either inflorescence in the binary test did not differ between treatments (post hoc Tukey test: control vs. decoy blue, odds ratio = 0.69 ± 0.38, z = -0.68, *p* = 0.77; control vs. decoy purple, odds ratio = 0.53 ± 0.30, z = -1.10, *p* = 0.51; decoy blue vs. decoy purple, odds ratio = 0.78 ± 0.43, z = -0.45, *p* = 0.89; **Fig. 3**) or within treatments (post hoc Wald *z*-test: control: log-odds = -0.50 ± 0.46, z = -1.07, *p* = 0.28; decoy blue: log-odds = -0.12 ± 0.47, z = -0.26, *p* = 0.79; decoy purple: log-odds = 0.13 ± 0.42, z = 0.31, *p* = 0.76).

**Figure 3:**
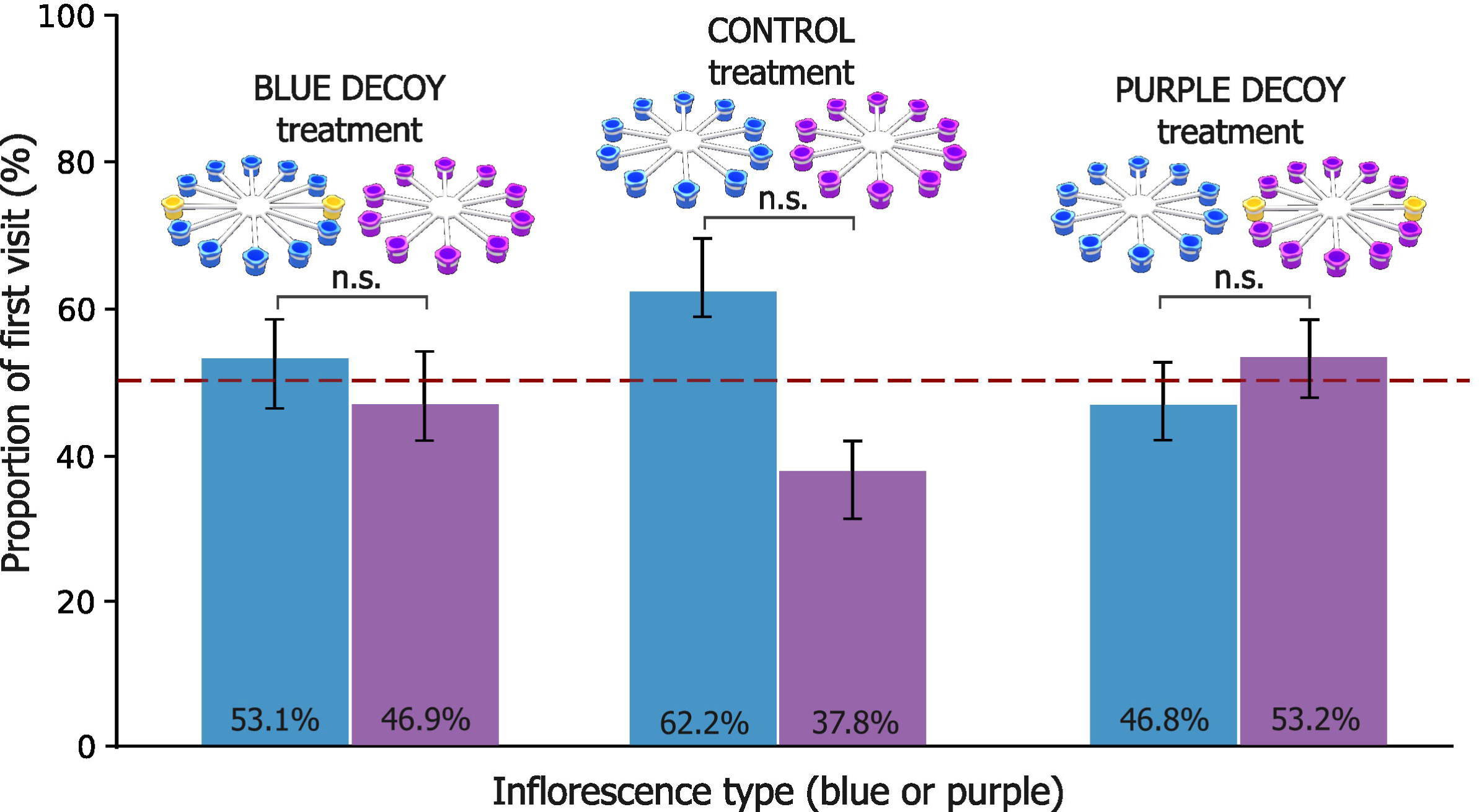
Estimated probabilities of first visits in the binary test across treatments. Probabilities were derived from the GLMM and calculated through post hoc analysis. Error bars represent the standard errors of the predictions. Statistical differences within treatments are indicated as n.s. (*p* > 0.05).

### Effect of probing a decoy flower on floral preferences

Probing an unrewarded flower did not affect bees’ preference between the blue and purple inflorescences in the next foraging bout (GLMM, binomial family; probed decoy: Χ² = 1.34, df = 5, *p* = 0.93). Preferences did not differ across bouts (foraging bout: Χ² = 5.47, df = 9, *p* = 0.79), nor did the effect of probing a decoy vary with foraging bout (interaction: Χ² = 1.33, df = 5, *p* = 0.93). The likelihood of first visiting the decoy-equipped inflorescence was similar for bees that probed a decoy flower in the previous bout and those that did not (**Table 1**).

**Table 1:**
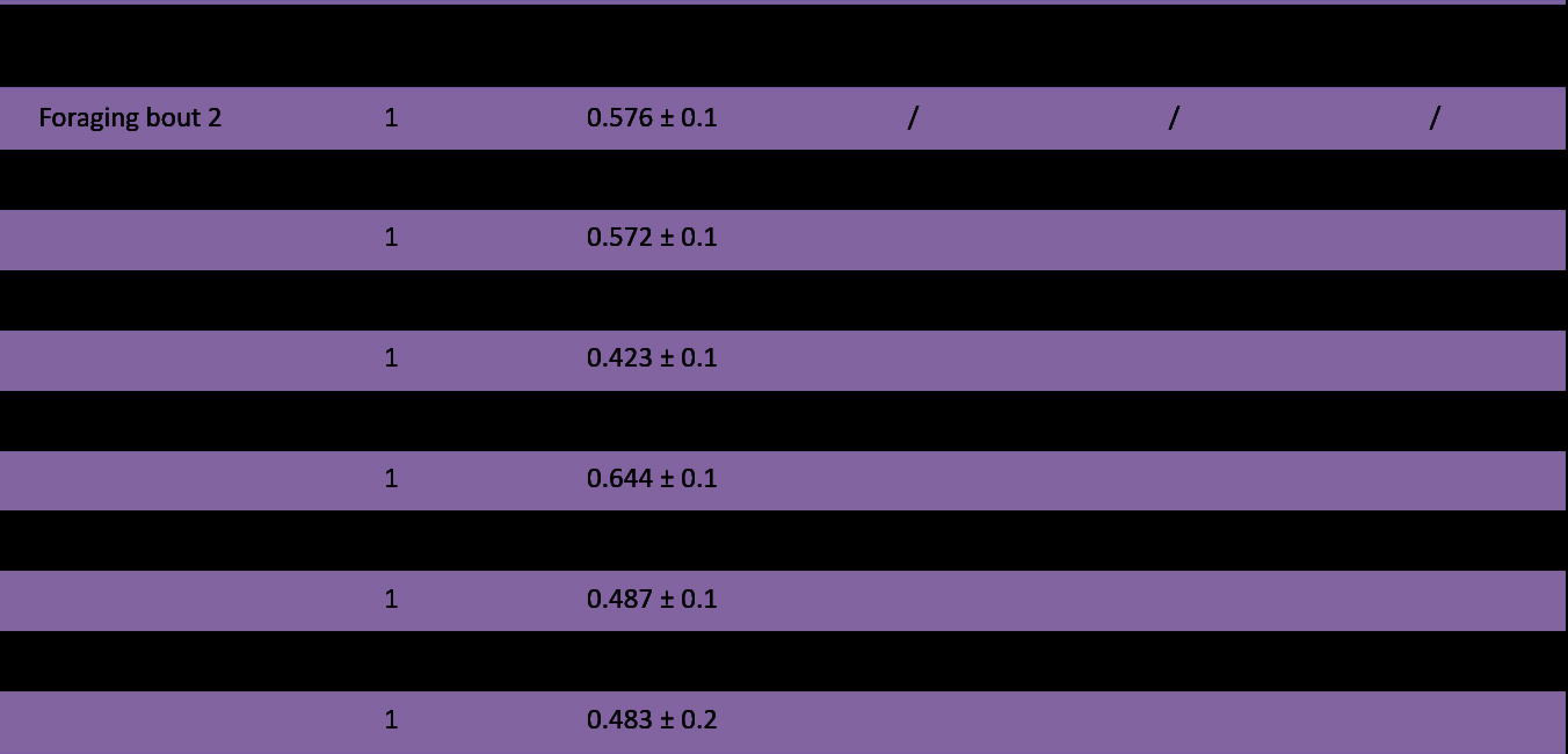
Estimated probabilities of first inflorescence choices and pairwise comparisons across bouts. Estimated probabilities are of first visits to the decoy-equipped inflorescences, based on the GLMM model. Odds ratios, 95% confidence intervals (CI), and p-values refer to pairwise comparisons between conditions (whether a decoy flower was probed in the previous bout or not).

### Frequency of interactions with decoy flowers across bouts

Bees quickly learned to avoid unrewarded flowers: foraging bout significantly affected the likelihood of probing decoy flowers (GLMM, binomial family; foraging bout: X² = 32.83, df = 4, *p* < 0.0001). Bees were less likely to probe unrewarded flowers in later bouts compared to earlier ones (**Fig. 4**). Additionally, of the 363 first flower landings recorded across all training bouts, 15 were on decoy flowers in the first bout (26% of landings in that bout), compared to just 2 across all subsequent bouts combined (0.7% of landings).

**Figure 4:**
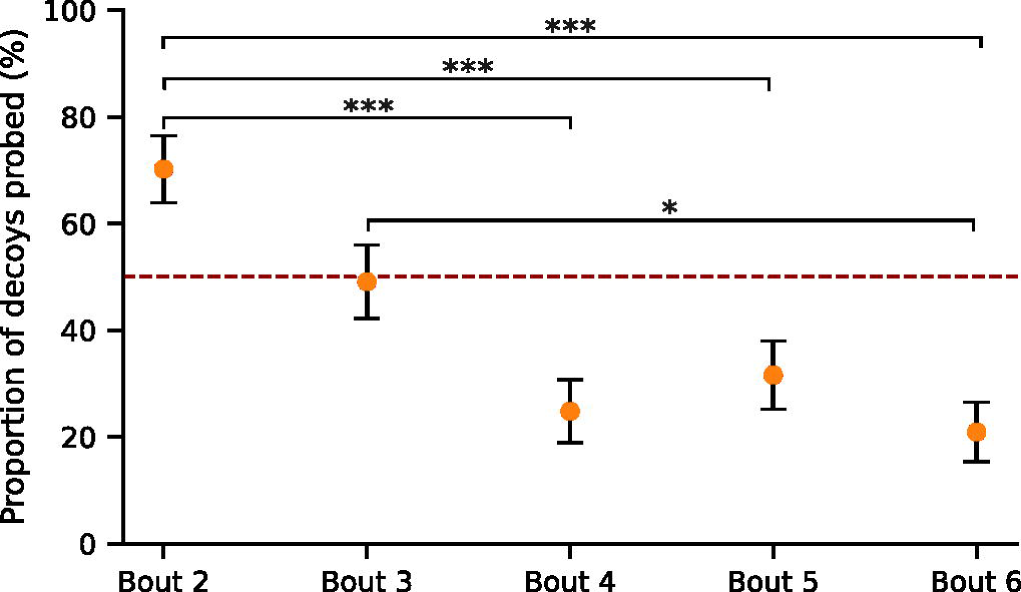
Estimated proportions of probing a decoy flower across foraging bouts. Probabilities of probing a decoy flower in training bouts 2–6 were derived from the GLMM and calculated through post hoc analysis. Error bars represent the standard errors of the predictions. Statistical differences within treatments are indicated as **\*\*\*** (*p* ≤ 0.001), and **\*** (p ≤ 0.05).

## DISCUSSION

We tested whether rewardless yellow “decoy” flowers influenced bumblebees’ choice between two equally rewarding inflorescences, blue and purple. Contrary to expectations, unrewarded flowers did not increase bees’ preference for neighbouring flowers. In the binary choice test, bees showed no overall preference for either inflorescence. During training, those that probed a rewardless flower did not favour rewarded flowers in the same inflorescence in the next bout. However, bees quickly learned to avoid unrewarded flowers, reducing visits and probings over successive bouts.

A decoy effect would have caused the introduction of an irrelevant option to shift bees’ relative preference between existing choices (Huber et al., 1982). In that case, first inflorescence choice in the binary test should have increased for the blue inflorescence in the “decoy blue” treatment and for the purple inflorescence in the “decoy purple” treatment, relative to the control (see **Fig. 3**). Our findings align with those of Forster et al. (2023a, b; 2025), who found that the presence of empty, previously high-reward flowers did not influence bees’ foraging choices. In their honeybee experiment (Forster et al., 2023a), decoy flowers did not shift preference between the two available flowers but led to increased movement between flowers and higher patch abandonment. In their study on stingless bees, Forster et al. (2023b) found no preference for the flower option matching the colour of the decoy flower. In Forster et al. (2025), bumblebee foraging choices were tested in a social context rather than individually, as in our study. They found that social information was the primary driver of choice, with bees preferring flowers where conspecifics were present.

During training, probing an unrewarded flower did not make bees more likely to favour neighbouring rewarded flowers in the next bout. Instead, their likelihood of choosing either inflorescence first remained constant across bouts. This contradicts our hypothesis that an irrelevant option would create an attraction effect, making nearby rewarding flowers more appealing by comparison (Huber et al., 1982), thereby increasing bees’ likelihood of foraging on these flowers rather than on similar ones in another inflorescence. However, it is important to note that rewardless flowers in our setup did not have a repulsion effect either: bees systematically collected all available sucrose rewards on an inflorescence before switching to the other, regardless of the presence of unrewarded flowers. By contrast, previous studies reported that rewardless flowers reduced pollinator visitation to an inflorescence (Cresswell, 1990; Smithson & Gigord, 2003; Forster et al., 2023b). Similarly, bumblebees were found to visit fewer flowers within an inflorescence when some were empty (Biernaskie et al., 2002; Smithson & Gigord, 2003; Nakamura & Kudo, 2016).

While unrewarded flowers did not influence foraging choices, bees in our experiment quickly learned to associate them with the absence of rewards. In decoy treatments, one-quarter of first landings were on unrewarded flowers in the initial bout, dropping to less than 1% over the next five training bouts. Bees also progressively reduced their probing of rewardless flowers across successive bouts. Extensive research showed that bees readily associate colours with rewards (Heinrich, 2004; Raine and Chittka, 2008) and recall spatial landmarks (Menzel et al., 1996), as seen in our setup where unrewarded yellow flowers remained in a fixed position across bouts. Similarly, studies have shown that bees quickly reduce visits to newly depleted flowers (Forster et al., 2023b; 2025), avoid unattractive food sources (Tan et al., 2014a), and minimise handling time of empty flowers (Smithson and Gigord, 2003).

In the control treatment without decoy flowers, bees tended to choose the blue inflorescence first more often than the purple one despite both being equally rewarding (see **Fig. 3**), but this preference was not significant. While bees have innate preferences for violet and blue flowers (Gumbert, 2000; Raine & Chittka, 2007), our purple flowers had lower spectral purity than the blue ones due to additional reflectance in the red range (see curves in **S2**), which likely reduced their saliency in the arena (Lunau, 1990). High spectral purity enhances floral attractiveness to bees (Lunau et al., 1996; Gumbert, 2000), which may explain the slight preference for blue. However, this tendency disappeared in the decoy treatments; we suggest that the presence of yellow flowers increased visual complexity, mitigating any subtle colour biases.

First visits to the yellow, unrewarded flowers were notably high (25%) during the first bout, despite their low abundance — only two (10%) among 20 blue and purple flowers in the arena. This may be due to their visual similarity to the white flowers used in pre-training, which bees had positively associated with rewards. Since the bees were colour-naïve in the initial bout, they may have perceived the yellow flowers as familiar. Bumblebees typically show a preference for novel colours that resemble those they have previously encountered (Gumbert, 2000).

While bees gradually reduced their probing of unrewarded flowers, they did not stop entirely. Instead, probings stabilised in later bouts rather than declining to zero, suggesting that bees continued to inspect them intermittently, albeit less frequently. Video analysis confirmed this pattern, showing that individual bees frequently re-probed rewardless flowers in a later bout even though they ignored them in previous ones. Similarly, Forster et al. (2023a) found that bees revisited empty flowers, possibly anticipating nectar replenishment. Ishii et al. (2008) noted that while bumblebees frequently re-inspected rewardless flowers, they re-probed them less often. In nature, bees commonly revisit depleted flowers probably due to memory constraints (Chittka et al., 1999).

Is it surprising that we found no clear decoy effect in our experiments, when they are apparently commonly reported? A closer look at the literature reveals that clear evidence for decoy effects is often lacking. For instance, studies testing asymmetrically dominated (AD) decoys in bees have failed to demonstrate attraction effects. In Shafir et al. (2002), where AD decoys were tested in honeybees and grey jays, only grey jays showed a true decoy effect. In bees, the decoy flower reduced choices for the competitor more than the target option, shifting relative preference, but this resulted from direct selection of the decoy rather than an increased attraction to the target. Similarly, Tan et al. (2014b) found that an AD decoy redirected choices from the competitor without increasing honeybees’ preference for the target food option. Hemingway et al. (2024) tested an AD decoy in bumblebees and found that adding a lower-concentration flower increased preference for the target, a medium-concentration flower. However, this shift likely resulted from an incentive contrast: rather than enhancing the perceived value of the medium option, exposure to the lower-quality alternative merely lowered the bees’ acceptance threshold. Incentive contrast effects are well-documented in bees, which often reject lower-quality rewards after experiencing higher-quality ones (Bitterman, 1976; Waldron et al., 2005; Townsend-Mehler et al., 2011). To our knowledge, the only example of a true decoy effect in insects was reported by Sasaki and Pratt (2011), who presented individual ants and whole colonies with two equivalent but different nest sites: one with a small entrance (preferred) but too bright, and the other dark (preferred) but with an overly large entrance. Individual ants shifted their preference toward the nest superior to the decoy nest they had previously experienced, whereas whole colonies showed no decoy effect. This is an important study, though we note the modest sample sizes of 12 and 14 ants per decoy treatment.

Likewise, previous studies testing phantom decoys in pollinators have been mostly inconclusive. Recent work by Forster et al. (2023a, 2023b, 2025) showed that empty flowers had no effect on preference or choice in several bee species. Tan et al. (2014a) found that while a few honeybees shifted toward the option most similar to an attractive phantom decoy, there was no consistent effect at the group level. The authors suggest this shift was due to incentive contrast, similarly to Hemingway et al. (2024): some bees that expected a high reward but found nothing lowered their selectivity and settled for a lower-quality alternative, rather than being drawn toward the most similar available option as predicted in a true decoy effect. Moreover, they found that an unattractive phantom decoy did not affect bee choice, instead of making the similar, still-available option look better in comparison and thus increase its selection rate.

The robustness of attraction effects has been increasingly questioned in recent years (Frederick et al., 2014; Yang and Lynn, 2014; Evans et al., 2021). Frederick et al. (2014) replicated 38 human studies on AD decoys but failed to reproduce most findings; they found attraction effects only when options were compared numerically, rather than experienced directly. The authors argued that in realistic settings, direct experience weakens structured comparisons, making dominance relationships less apparent. Additionally, attraction effects may diminish or disappear when decisions are made under time constraints, as often occurs in real-world situations (Pettibone, 2012; Marini et al., 2023). This issue is particularly relevant for nonhuman animals, where decoy effects can only be tested through direct choices. In fact, numerous studies on decoy effects in nonhuman animals had mixed or partial results (Bateson et al., 2003; Scarpi, 2011; Pinto et al., 2016; Marini et al., 2023), or found no evidence of a decoy effect at all (Bateson, 2002; Edwards and Pratt, 2009; Cohen and Santos, 2017; Parrish et al., 2018). To our knowledge, there is currently no clear demonstration of a decoy effect in pollinators.

Could unrewarded flowers induce repulsion effects on neighbouring flowers instead? Flower-visiting insects have been shown to visit fewer flowers within inflorescences containing empty flowers (Biernaskie et al., 2002; Ishii et al., 2008), or even avoid patches entirely when rewardless flowers are abundant (Hirabayashi et al., 2006; Nakamura and Kudo, 2016). Notably, some decoy studies have reported repulsion effects, suggesting that a decoy option can make the target appear less desirable, as the negative perception of the decoy “contaminates” the target (Frederick et al., 2014). Spektor et al. (2018) found that such repulsion effects emerge when the decoy and target options are closely located, with their proximity triggering a similarity effect between them.

Despite their potential repulsive effects, a balanced proportion of nectarless flowers may be beneficial to plants (Bell, 1986). Bees prefer larger inflorescences over smaller ones (Hirabayashi et al., 2006; Ishii et al., 2008), so bearing rewardless, “cost-free” additional flowers may enhance the attractiveness of an inflorescence (although we found no evidence for this in our study). Moreover, fewer visits per inflorescence due to the presence of nectarless flowers can help reduce selfing, limiting inbreeding and thereby, enhancing plant fitness (de Jong et al., 1993; Biernaskie et al., 2002).

In summary, we found that the addition of rewardless flowers did not increase bees’ preference for nearby rewarded flowers. While not a classical asymmetrically dominated decoy study, the experiment shared many elements with such studies, and has direct ecological relevance to rewardless flowers on inflorescences. Bees rapidly adapted to the presence of rewardless flowers, reducing the time spent visiting and probing them after just a few foraging bouts. A sizeable body of research is accumulating, suggesting that classical decoy effects do not play a significant role in insect foraging, and thus decoy-like effects cannot explain the presence of nectarless flowers. Further research is needed to clarify the effect of nectarless flowers on the attractiveness of surrounding flowers. While decoy and decoy-like effects attract a lot of research attention, evidence for them is weak and inconsistent, and it may be time to redirect research attention to other valuation illusions.

## SUPPLEMENTARY MATERIAL

**S1** contains the statistical analysis of the main experiment, **S2** provides the reflectance curves of the artificial inflorescences, **S3** includes a video clip of a bee foraging during the experiment, and **S4** the dataset. All supplementary materials are available on Zenodo (https://doi.org/10.5281/zenodo.14995342).

## ACKNOWLEDGMENTS

We want to thank B. Rubene and L. Romrig for their assistance with data collection. Special thanks to A. Avarguès-Weber for providing spectrometer measurements of the artificial flowers.

## AUTHORS’ CONTRIBUTION

**Mélissa Armand:** Conceptualization, Methodology, Software, Validation, Formal analysis, Investigation, Writing - Original Draft, Writing - Review & Editing, Visualization. **Leonhard Herrnberger:** Investigation, Methodology. **Clara Jung:** Investigation. **Tomer J. Czaczkes:** Conceptualization, Methodology, Validation, Resources, Writing - Review & Editing, Supervision, Project administration, Funding acquisition.

